# BPIFB1 loss alters airway mucus properties and diminishes mucociliary clearance

**DOI:** 10.1101/2020.06.17.155846

**Authors:** Lauren J. Donoghue, Matthew R. Markovetz, Cameron B. Morrison, Kathryn M. McFadden, Troy D. Rogers, Brian Button, Camille Ehre, David B. Hill, Barbara R. Grubb, Samir N. P. Kelada

**Affiliations:** Department of Genetics, University of North Carolina at Chapel Hill; Curriculum in Genetics and Molecular Biology, University of North Carolina at Chapel Hill; Marsico Lung Institute, University of North Carolina at Chapel Hill; Department of Biochemistry and Biophysics, University of North Carolina at Chapel Hill; Division of Pediatric Pulmonology, University of North Carolina at Chapel Hill; Department of Physics and Astronomy, University of North Carolina at Chapel Hill

## Abstract

Airway mucociliary clearance (MCC) is required for host defense and often diminished in chronic lung diseases. Effective clearance depends upon coordinated actions of the airway epithelium and a mobile mucus layer. Dysregulation of the primary secreted airway mucin proteins, MUC5B and MUC5AC, is associated with a reduction in the rate of MCC; however, how other secreted proteins impact the integrity of the mucus layer and MCC remains unclear. We previously identified the gene *Bpifb1/Lplunc1* as a regulator of airway MUC5B levels using genetic approaches. Here, we show that BPIFB1 is required for normal mucociliary clearance *in vivo* using *Bpifb1* knockout (KO) mice. Reduced MCC in *Bpifb1* KO mice occurred in the absence of defects in sodium or chloride ion transport or reduced ciliary beat frequency. BPIFB1 loss resulted in airway mucus flakes with significantly increased complex viscosity, a key biophysical property of mucus known to impact MCC. Finally, we detected colocalization of BPIFB1 and MUC5B in secretory granules in mice and in the protein mesh of secreted mucus in human airway cultures. Collectively, our findings demonstrate that BPIFB1 is an important component of the mucociliary apparatus in mice and a key component of the mucus protein network.

## INTRODUCTION

The airway mucus layer plays a critical role in airway homeostasis and host defense in part by facilitating mucociliary clearance (MCC) of inhaled particles and pathogens. Effective MCC requires a properly hydrated airway surface, an optimal balance of components in secreted mucus (i.e., proteins, salt), and coordinated beating of cilia. The importance of the MCC apparatus is highlighted by the fact that mutations in genes essential to ciliary function or ion transport cause primary ciliary dyskinesia (PCD) and cystic fibrosis (CF), respectively, in which respiratory infections are prominent and severe. In the context of more common obstructive airway disease such as asthma and chronic obstructive pulmonary disease (COPD), changes in MCC have been associated with variation in the abundance of the secreted mucin proteins MUC5B and MUC5AC, which are key contributors to the gel-like nature of healthy airway mucus^1^. The concentration, organization, and post-translational modifications of MUC5B and MUC5AC also have been shown to alter biophysical properties of mucus that are required for effective MCC^2–6^. Furthermore, the ability of mucins to form a proper gel-structure is dependent upon interactions with other secreted proteins, referred to as the “mucin interactome”^7^, although the specifics of these dependencies remain unclear.

In contrast to MUC5AC, which is typically only observed at high levels in disease states, MUC5B is constitutively expressed in the airways of healthy individuals where it has a role in regulating homeostasis through airway MCC and host defense^8,9^. We previously identified the gene *Bpifb1* (also known as *Lplunc1*) as a regulator of airway MUC5B protein levels using a quantitative genetics approach in mice and showed that loss of BPIFB1 in a knockout (KO) mouse model led to increased levels of MUC5B protein in the airways^10^. Given that MUC5B levels have been shown to affect MCC^9,11^, we sought to determine if *Bpifb1* also had a role in regulating MCC. We observed a substantial loss of MCC in the lower airways of *Bpifb1* KO mice. To identify which aspects of the MCC apparatus contributed to this defect, we assessed epithelial sodium and chloride ion transport, ciliary function, and biophysical properties of mucus. Our results show that alterations in the mucus layer including increased mucus viscoelasticity were associated with decreased MCC in the absence of BPIFB1 and that BPIFB1 is a component of the mucus protein network.

## RESULTS AND DISCUSSION

In order to determine if *Bpifb1* has a role in MCC, we assessed MCC in tracheas of naïve *Bpifb1* KO and WT mice. We quantified MCC by measuring the percentage of instilled beads that were cleared by the trachea over a fixed period of time. *Bpifb1* KO mice exhibited significantly decreased clearance, with some mice exhibiting no tracheal bead clearance whatsoever (Figure 1A). These results indicated clearly that BPIFB1 is required for MCC.

**Figure 1.**
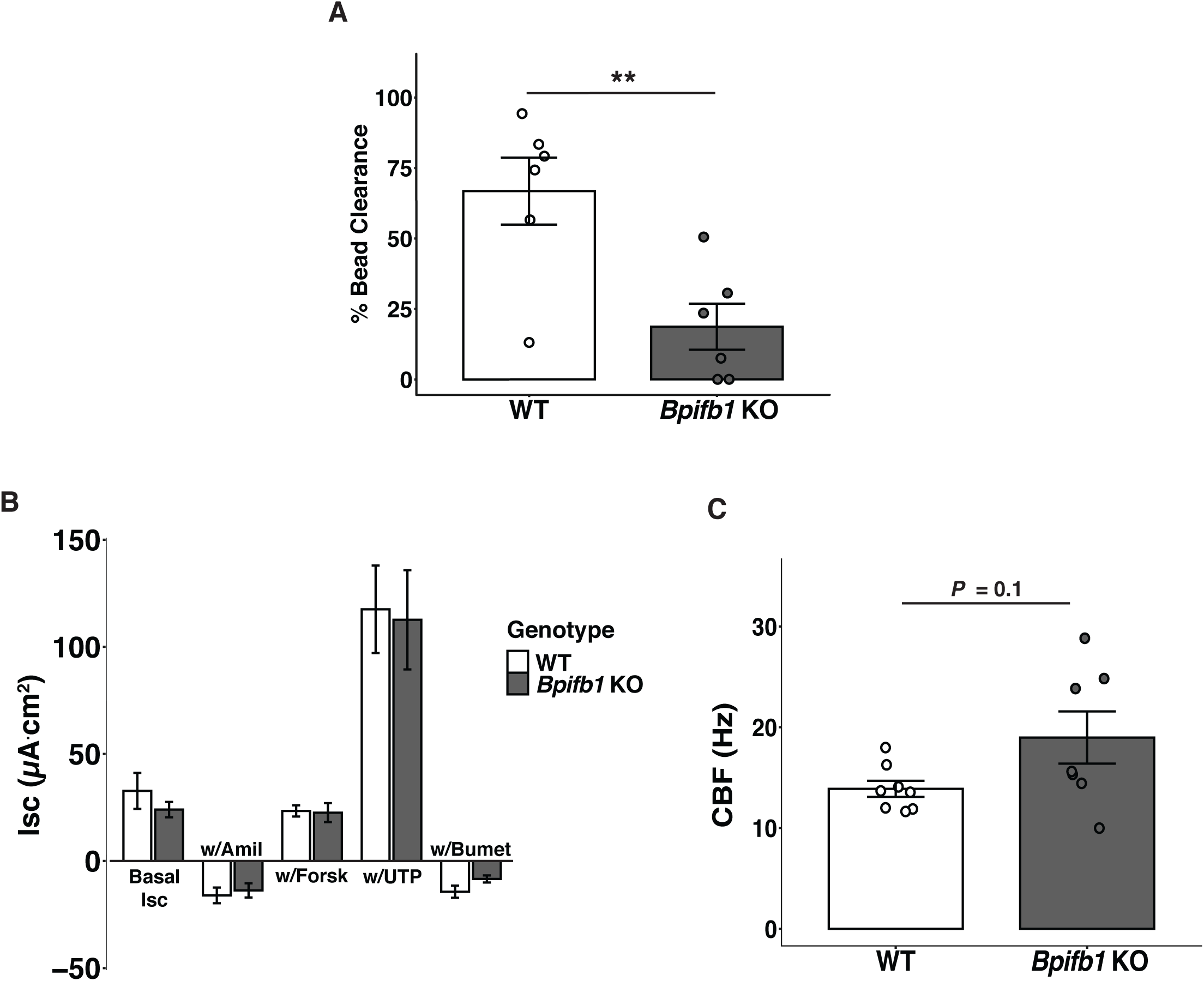
Decreased mucociliary clearance in *Bpifb1* KO mice. (A) Percent bead clearance in tracheal mucociliary clearance assay in naive WT and *Bpifb1* KO mice. Mean ± SEM depicted. n=6/genotype. Significance of two-sided Student’s *t*-test denoted as **, *P* < 0.01. (B) Ion transport measured in Ussing chambers. Mean ± SEM depicted. n=6/genotype. No differences between genotypes by two-sided Student’s *t*-test. (C) Ciliary beat frequency by *in situ* tracheal assay. Mean ± SEM depicted. n=7-8/genotype. Difference between genotypes not significant by two-sided Welch’s *t*-test, *P* = 0.1.

To understand the mechanism by which BPIFB1 loss leads to defective MCC, we systematically tested the expression and function of key components of the MCC apparatus. First, we examined the expression of key ion transporters in the airway epithelium given their importance in regulating airway hydration and MCC^1^. RNA sequencing of lung tissue from *Bpifb1* KO and WT mice did not reveal significant differences in expression of well-characterized sodium and chloride ion transporter genes including *Cftr, Scnn1b*, and *Tmem16a* (Figure S1). However, given that a lack of difference in ion transporter gene expression does not rule out a difference in functionality, we also measured bioelectric capacities in *Bpifb1* KO and WT tracheas. We found no differences in short-circuit current (*I*_*sc*_) at baseline or following stimulation with compounds targeting the major ion channels in the airway epithelium (i.e. CFTR, ENaC, CaCC), (Figure 1B). This indicated that the loss of MCC in *Bpifb1* KO mice was likely not driven by altered ion transport. Additionally, as ciliary beating is essential for MCC, we directly measured ciliary beat frequency (CBF) in intact tracheas *in situ*. In comparison to WT mice, *Bpifb1* KO mice had a slightly elevated CBF, albeit not statistically significant at *P* < 0.05 (Fig 1C). These data indicated that reduced MCC was not associated with reduced ciliary beating.

Next, to determine if *Bpifb1* loss influenced the expression of other genes or biological processes that could underlie the defect in MCC, we examined global gene expression in lung tissue from *Bpifb1* KO and WT mice. Excluding *Bpifb1*, only 4 protein coding genes and 2 long non-coding RNAs were differentially expressed by genotype (|fold change| ≥ 2 and adjusted *P*-value ≤ 0.05, Figure S1). Of these genes, none have been previously annotated for a direct role in MCC or related pathways. Furthermore, genes related to mucin production, including *Muc5b, Muc5ac, Spdef*, and *Foxa3*, were not differentially expressed. These gene expression results suggested the reduction in MCC in *Bpifb1* KO mice was not mediated by differential expression of key MCC or mucus production genes.

Given the lack of differences in hallmark causes of reduced MCC, we investigated if there were differences in the airway mucus *per se* that could be contributing to the observed loss of MCC. Using microbead rheology, we measured the complex viscosity (η*), a measure of both viscosity and elasticity, of mucus flakes (i.e. the insoluble portions of airway mucus) in bronchoalveolar lavage (BAL) fluid from *Bpifb1* KO vs WT mice. For these measurements, mice were intranasally exposed to house dust mite allergen for three weeks to increase the levels of mucus in BAL, and there were no differences in the degree of eosinophilic inflammation by genotype (Supplementary Figure S2). We found that the complex viscosity of mucus flakes was significantly increased in *Bpifb1* KO mice compared to WT mice (Figure 2A, B). By imaging of whole BAL, we determined this increase was independent of the size and number of mucus flakes, and there were no differences in these metrics by genotype (Supplementary Figure S3A-E). Consistent with these findings and our previous report of increased MUC5B in BAL from naïve *Bpifb1* KO mice^10^, we found that MUC5B levels were elevated by genotype in the supernatant fraction of centrifuged BAL from naïve mice, but not in the pelleted (i.e. insoluble) fraction (Figure 2C). Thus, loss of BPIFB1 results in altered biophysical properties of mucus flakes and increased levels of soluble MUC5B.

**Figure 2.**
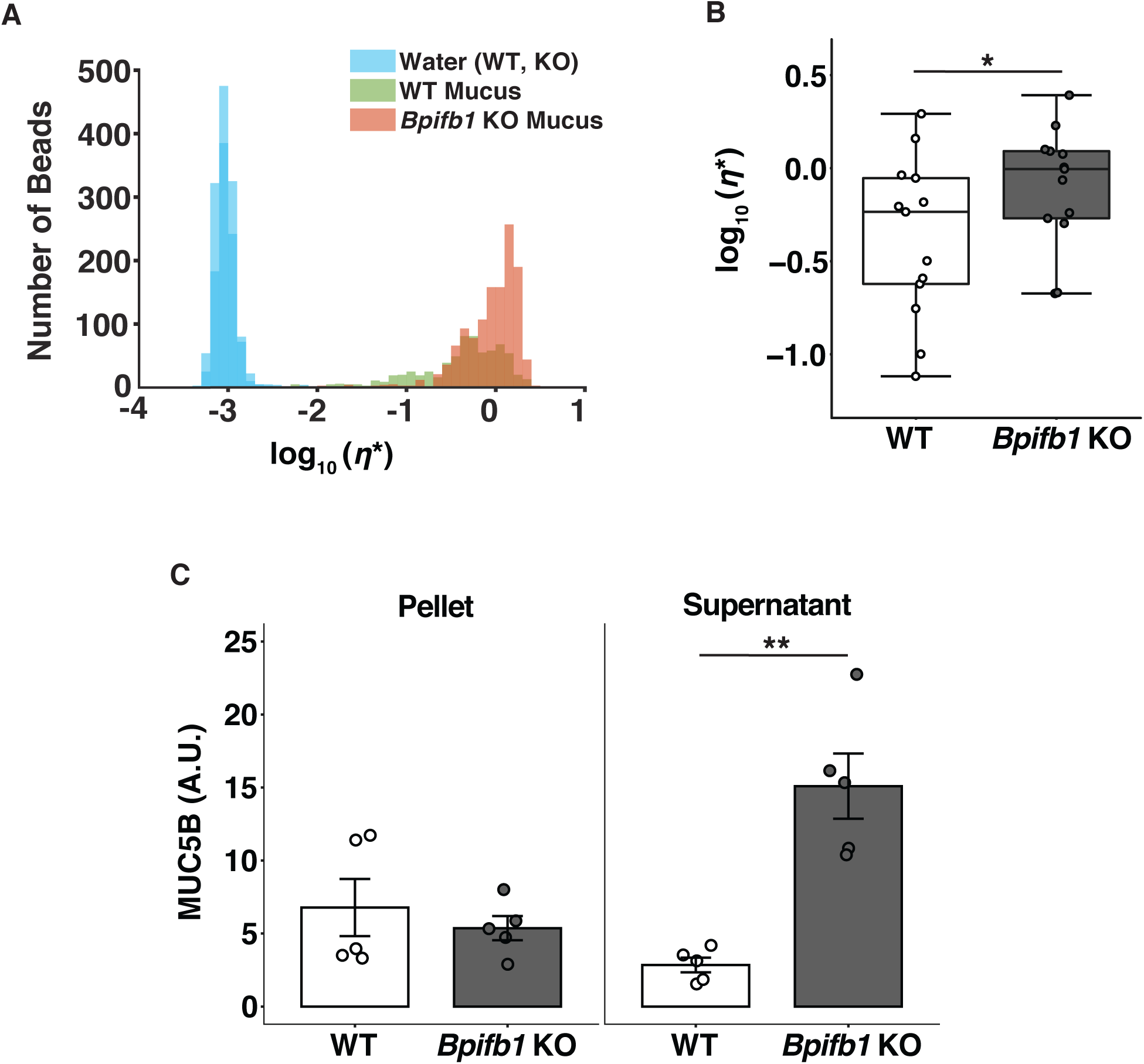
Biophysical properties of airway mucus are altered in *Bpifb1* KO mice. (A) Distribution of rheological signal in BAL from a representative allergen exposed WT and *Bpifb1* KO mouse. (B) Complex viscosity of BAL mucus fraction. Boxplots depict median and interquartile ranges. n=13/genotype. Significant difference between genotypes by ANOVA modeling denoted as *, *P* < 0.05. (C) MUC5B in pellet and supernatant fractions of BAL from naive WT and *Bpifb1* KO mice by mucin western blotting. Mean ± SEM depicted. Significant difference between genotypes in supernatant fraction by two-sided Welch’s *t*-test denoted as **, *P* < 0.01. n=5/genotype.

To gain insight into whether BPIFB1 colocalizes with mucin protein in the airways and, if so, whether this association occurs before mucin granule release, we assessed the spatial relationship between BPIFB1 and mucin protein using fluorescent immunostaining. By colocalization analysis of labeled BPIFB1 and MUC5B protein in large airways of mice challenged with allergen, we determined that the vast majority of MUC5B positive cells were also positive for BPIFB1 (Figure 3A, B), indicating regulated cellular co-expression of MUC5B and BPIFB1. Colocalization was also present in the airways of mice not challenged with allergen (Supplementary Figure S4). Furthermore, MUC5B positive granules were also positive for BPIFB1 and luminal mucus contained both MUC5B and BPIFB1 (Figure 3C), suggesting that packaging and secretion of these proteins is likely coupled.

**Figure 3:**
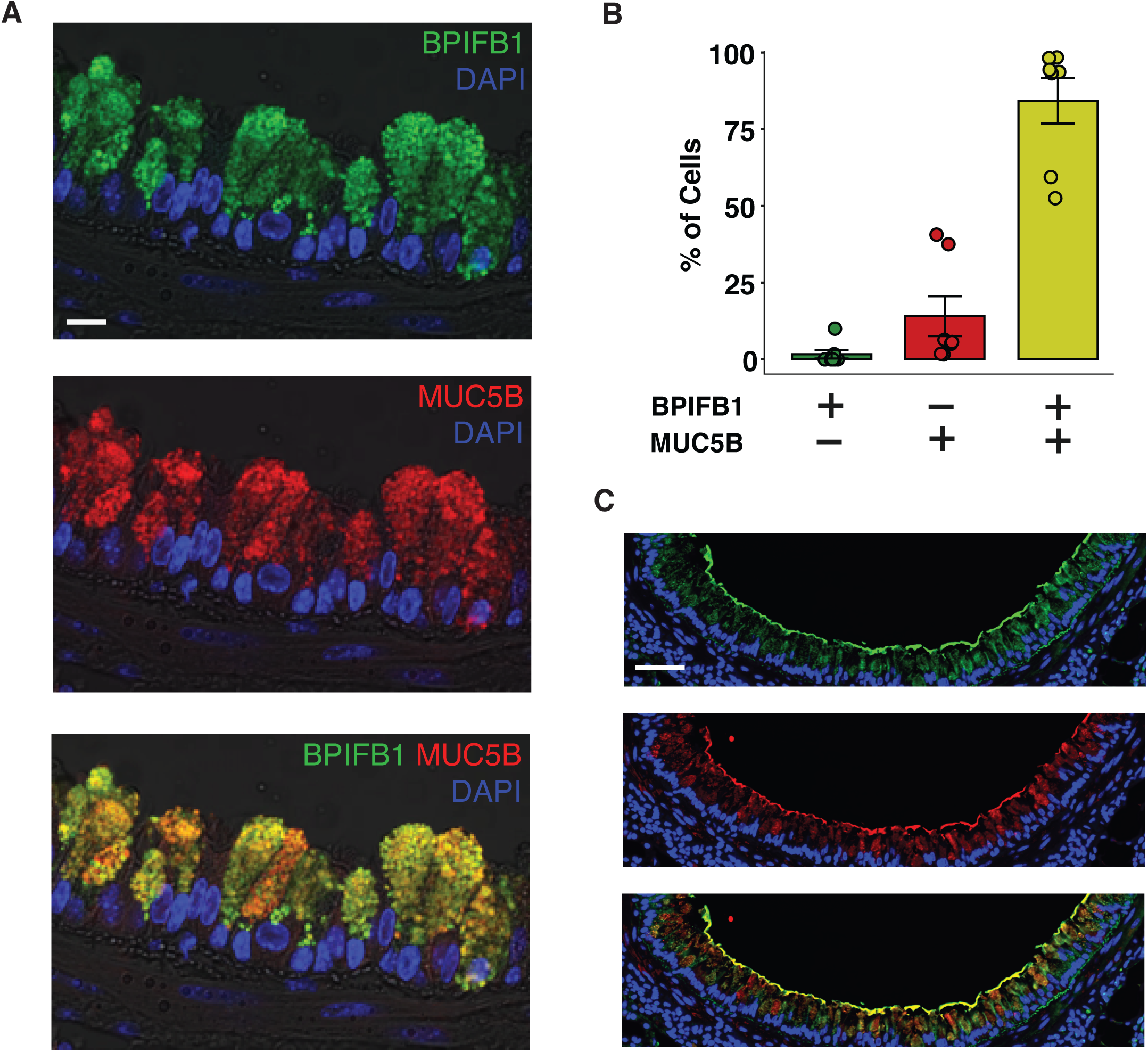
BPIFB1 and MUC5B colocalize in murine airway epithelial cells. (A) BPIFB1 (green) and MUC5B (red) immunostaining in airway epithelial cells from allergen challegned mice. Scale bar = 10 *µ*m. (B) Percentage of cells positively stained for BPIFB1 only, MUC5B only, or both in allergen challenged mice. n=7 mice. (C) BPIFB1 (green) and MUC5B (red) immunostaining in large airways of allergen challegned mice demonstrating BPIFB1 staining in luminal mucus. Scale bar = 50 *µ*m.

Finally, we performed immunogold-labeled scanning electron microscopy for BPIFB1 and MUC5B in human bronchial epithelial cells (HBECs) cultured at air-liquid-interface to determine if BPIFB1 closely associated with extracellular mucus. We detected BPIFB1 exclusively in mucus strands coating the cellular surface (Figure 4). This finding confirmed that BPIFB1 is a component of the mucus protein network produced by human airway cells.

**Figure 4:**
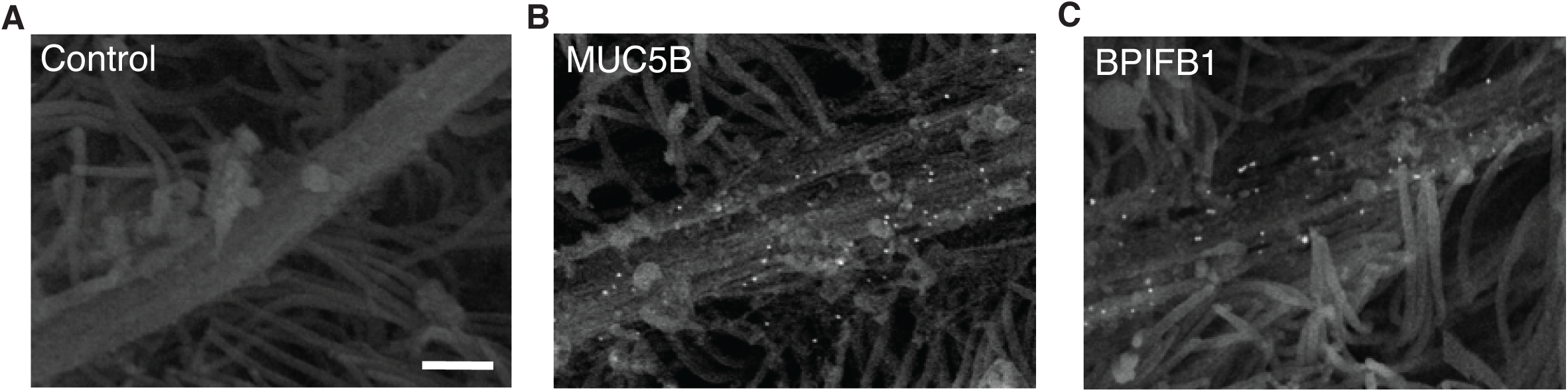
BPIFB1 is part of mucus netwrok in human bronchial epithelial cell culutures. Immunogold-labeled scanned electron microscopy for (A) negative control, (B) MUC5B, and (C) BPIFB1. Scale bar = 1 *µ*m.

Significantly altered levels of either *BPIFB1* gene or protein expression have been observed in CF, COPD, asthma, and IPF^12–15^, indicating a strong connection between *BPIFB1* and muco-obstructive diseases. After identifying *Bpifb1* as a regulator of airway mucus levels through an unbiased genome-wide search in the mouse^10^, we sought to interrogate the specific function of BPIFB1 in the airways. We found that the loss of BPIFB1 reduced MCC dramatically in naïve *Bpifb1* KO mice, demonstrating the critical role of this gene in regulating MCC. Thus, in addition to the strong connection between *BPIFB1* gene/protein expression and muco-obstructive diseases, our findings indicate BPIFB1 may have an important role in airway host defense by regulating MCC. Studies in humans have shown that there is significant interindividual variation in baseline MCC, and analysis of MCC in monozygotic and dizygotic twins has implicated genetic variation as a modifier of MCC^16^. Given the identification of common genetic variation associated with the levels of *BPIFB1* gene expression^17^, genetic variation in *BPIFB1* regulation could be one such factor regulating baseline MCC in humans. Furthermore, BPIFB1 was one of the top 20 proteins detected in endotracheal tube mucus from individuals without a history of lung disease or smoking, highlighting its abundance in non-disease states^18^.

Due to the lack of differences in basal or stimulated transport of sodium and chloride ions, ciliary beating, and MCC-related gene expression between *Bpfib1* KO and WT mice, our data point toward changes to the airway mucus layer as the cause of reduced MCC. More specifically, the increased complex viscosity of mucus, as we observed in mucus flakes from *Bpifb1* KO mice and may be the case for soluble mucus as well, is suggestive of mucus that is more difficult to clear from the airways. *In vitro* studies with human airway cultures^19–22^, as well as earlier studies in model organisms^23–25^, have demonstrated that mucus transport is directly related to the rheological properties of mucus. The increased CBF that we observed in *Bpifb1* KO mice, although not statistically significant, may be indicative of an attempt to compensate for altered airways mucus^26^. Such a phenomenon has been demonstrated in human bronchial epithelial cultures where CBF increased following a moderate increase in mucus viscosity or mucus load^27^.

The increased complex viscosity of mucus flakes in *Bpifb1* KO mice may be attributable to a change in mucin network structure, which is a function of the density and/or nature of mucin-mucin/mucin-protein interactions^7^. In muco-obstructive airway disease contexts, for example, heightened mucus viscoelasticity due to oxidation-dependent mucin cross-linking^5^ or increased mucin concentration^28^ has been described. Direct measurement of mucin concentration was not possible in our experiments due to the high dilution factor of mucus in lavage fluid. However, the lack of increased MUC5B in pelleted BAL from *Bpifb1* KO mice provides some evidence against the hypothesis that mucin concentration is increased in mucus flakes and is the driver of changes in mucus rheology. That said, we detected elevated levels of soluble MUC5B in *Bpifb1* KO mice, which raises the question of whether MUC5B levels are higher in *Bpifb1* KO mice simply due to decreased MCC or increased mucin concentration. *In vitro* studies, in which mucus can be collected directly from the surface of epithelial cells, will be required to determine specifically how BPIFB1 impacts the mucus layer.

Although the polymeric nature of gel-forming mucins is essential for the biophysical properties of mucus, studies have shown that mucins alone cannot create complex networks, and thus, other factors, including secreted globular proteins, are necessary for mucus formation^29^. Our detection of BPIFB1 in secreted mucus from HBECs confirms BPFIB1 is indeed a component of the mucus protein network and supports the conclusion that this non-mucin protein plays an important role in the mucus layer. Further evidence for a direct role of BPIFB1 in the mucus network comes from a finding that BPIFB1 has strong physical interactions with mucin proteins in HEBC secretions. *Radicioni et al*. found that after treating mucus from HBEC cultures with guanidinium hydrochloride and detergents (i.e. dissociative conditions), BPIFB1 was one of few globular proteins interacting with mucins^7^. Additionally, our data demonstrating shared regulatory mechanisms between BPIFB1 and MUC5B suggest strongly coordinated roles. We previously found that *Bpifb1* and *Muc5b* gene expression was strongly correlated in mouse lung tissue^10^, and here we have shown that BPIFB1 and MUC5B protein are colocalized in airway epithelial cells and present in secreted mucus. Cellular co-expression between *BPIFB1* and *MUC5B* mRNA has also been demonstrated in human airways by single cell RNA sequencing studies^30,31^ and between BPIFB1 and MUC5AC protein in COPD^32^. Lastly, our finding that BPIFB1 is present in mucin secretory granules, which are tightly packed and only contain the necessary proteins for mucus, implicates an additional regulatory mechanism coupling these two proteins. Whether there is an optimal stoichiometry between levels of secreted BPIFB1 and mucin proteins for MCC, as may be the case between MUC5B and MUC5AC^2,33^, remains to be determined.

In summary, we demonstrate that BPIFB1 is critical for MCC *in vivo*. These findings establish that mucociliary dysfunction can be driven by non-mucin protein components of airway mucus, highlighting their importance in health and disease.

## METHODS

A detailed descriptions of the methods used in this study can be found in the Supplementary Methods (Supplementary File 1).

### Study Approval

All experiments conducted with mice in this study were compliant with an Institutional Animal Care and Use Committee protocol at an animal facility approved and accredited by the Association for Assessment and Accreditation of Laboratory Animal Care International. Primary cells were provided by the Cystic Fibrosis (CF) Center Tissue Core Facility of the University of North Carolina at Chapel Hill under the auspices of protocols approved by the Institutional Committee on the protection of the rights of human subjects.

### Data availability

Complete raw data and counts generated by RNA sequencing were deposited in the NCBI’s Gene Expression Omnibus (GEO) database under entry GSE150452.

## AUTHOR CONTRIBUTIONS

LJD, MRM, CBM, BB, CE, DBH, BRG, and SNPK conceived experiments and interpreted data. LJD, MRM, CBM, KMM, TDR, CE, and BRG performed experiments. LJD and SNPK wrote and revised the manuscript. All authors edited and reviewed the manuscript.

## ACKNOWLEDGEMENTS

We thank Aiman Abzhanova for assistance with immunostaining. This work was supported in part by NIH HL122711 (SNPK). LJD was supported by NIH F31HL143853 and a grant from the National Institute of General Medical Sciences under award 5T32 GM007092.

## FIGURE LEGENDS

Supplementary Figure Legends provided in Supplementary File 1.

