## Supplementary File 1 for "BPIFB1 loss alters airway mucus properties and diminishes mucociliary clearance"

**Table of Contents**

Supplemental Methods………………………….……………2-5
Supplementary Figure Legends………………….…………6
Supplementary Figures 1-4……………………………….7-10

**SUPPLEMENTAL METHODS**

*Mice and bronchoalveolar lavage.*

*Bpifb1* KO mice were generated as previously described[10]. Female and male homozygous WT and KO littermates were used in all experiments. Investigators were blinded to mouse genotype during all data collection and quantification, and sample order was randomized with respect to genotype. All mice were used at 8-12 weeks and naïve unless otherwise noted. For rheology experiments, mice were intranasally exposed to 25 μg house dust mite allergen (Stallergenes Greer, Cambridge, MA) three times per week for three weeks. Seventy-two hours following the last exposure, bronchoalveolar lavage (BAL) was performed by instilling 500uL of PBS with protease inhibitors into the lungs twice with the same fraction. For differential cell counts on mice under the same allergen protocol, an additional cohort of mice were lavaged with 500uL and 1mL fractions of PBS with protease inhibitors. BAL was spin at 500g for 10 min to pellet cells, cells were spun onto slides, and stained with Shandon Kwik Diff Stains (Thermo Scientific). Inflammatory cell counts by two investigators were averaged.

*Mucociliary clearance.*

MCC was measured using a previously described method[34]. Briefly, mice were anesthetized and fluorescent beads were inserted in 200nL PBS through a small incision to approximately the main stem bronchi. After 15 min, tracheas were removed, the tissue was dissolved, and the number of remaining beads were counted.

*RNA isolation, sequencing, and differential expression analysis.*

Lung tissue was harvested from naïve *Bpifb1* WT and KO mice and placed in RNA*later* (Ambion, MilliporeSigma, St. Louis, MO) for 2 weeks. RNA from the right middle lobe was isolated using an RNeasy Kit (Qiagen, Valencia, CA). Sequencing libraries were prepared by the UNC High Throughput Sequencing Facility using the KAPA stranded RNA-seq kit (Roche, Wilmington, MA) and sequenced on a HiSeq4000 (Illumina, San Diego, CA) to a depth of 20 million 50bp single-end reads/sample. Raw reads were evaluated with FastQC (http://www.bioinformatics.babraham.ac.uk/projects/fastqc) and MultiQC[35]. Reads were aligned to the C57BL/6J genome (GENCODE, release M18) using STAR version 2.6.0[36] and quantified with Salmon version 0.9.1[37]. Differential expression was performed using DESeq2 version 3.0 with the res() function with contrast KO vs WT[38]. Transcripts with fewer than 10 reads in more than 5 samples were excluded from analysis. Complete differential expression results are provided in Supplementary File 2.

*Ciliary beat frequency.*

Ciliary beat frequency (CBF) was measured in situ in the closed trachea, immediately after exsanguination of the mouse anesthetized with 3% isoflurane. For the tracheal preparation, the skin was opened, the muscle and connective tissue overlying the trachea were retracted and the exposed trachea was covered with a piece of plastic wrap dipped in water equilibrated mineral oil. This preparation was immediately placed under a dissecting scope (10x magnification) outfitted with a digital camera (Basler, Germany) and recorded with Basler software. The preparation was lighted a red LED  allowing light reflected from the beating cilia to be easily seen. The temperature the preparation was closely monitored using a tiny temperature probe (T Type insect probe, Physitemp Inst. Clifton N. J) placed alongside the trachea. The output from the temperature probe was displayed digitally on a Physitemp TCAT-2ac Controller and a small ceramic heater (Wuhostam, China) positioned close to the preparation was used to maintain the temperature of the preparation at 37 ($\pm$ 0.1°C ). We found it necessary to place the preparation on an air table (TMC, Peabody MA) to minimize vibrations that interfered with the CBF measurements. It took approximately 3 min from the time of euthanasia until the preparation was under the scope and data acquisition commenced. CBF was measured for a 2 s period at 10, 15, and 20 min after euthanasia. The data were collected at 100 frames/second and analyzed using SAVA software. CBF is reported as the mean of values from the three time points for each mouse. Data from one mouse was excluded because data were missing for two of the three time points.

*Ussing chamber bioelectrics.*
The bioelectrical proprieties of excised mouse tracheas following euthanasia were obtained using Ussing chambers as previously described [34]. Krebs-Ringer bicarbonate solution was added to the tissue and baseline measurements were taken (Isc), followed by the addition of amiloride ($10^{-4}$ M), forskolin ($10^{-5}$ M), UTP ($10^{-4}$ M), and Bumetanide ($10^{-4}$M) added sequentially to the basolateral side to block $Cl^-$ secretion.

*Microbead rheology.*
Whole/raw BAL samples from allergen exposed mice were mixed with 1-μm carboxylated microspheres. The diffusion of beads through the mixture was tracked and the displacement was estimated using previously described methods[9]. Complex viscosity of the insoluble mucus (flake) fraction of samples was determined using a Gaussian mixture model as described in Esther et al. Samples were excluded from the analysis if their complex viscosity was less than or equal to -2.7, indicating not enough mucus was present in the sample for an accurate measurement.

*Whole BAL Imaging.*
Whole/raw BAL samples from allergen exposed mice used for microbead rheology measurements were used for imaging. Tiled images were obtained by stitching together sequentially captured viewing fields at 40x in 11 BAL samples per genotype with 3 slides containing 5uL of BAL imaged in each sample. For each field both a fluorescence and a phase contrast image were captured with 10ms exposures in both modalities on an Olympus IX70 inverted light microscope. Images were segmented based on fluorescence intensity into flake and non-flake regions semi-automatically using a MATLAB program (The MathWorks, 2017) and single, user-specified threshold parameter. The threshold that provided the best user-adjudged outline of flake matter, also taking the phase contrast image into account, was used for final determination of flake regions. High-intensity regions of area < 500px (≈25μm in diameter) were excluded as too small to be distinguished from cells or cell debris.

*Mucin immunoblotting.*

Naïve mice were lavaged with 500uL PBS with protease inhibitors and lavage was spun at 4,000 rpm for 10 min at 4°C. The lavage was then separated into a supernatant fraction and pellet fraction. The pellet fraction was resuspended in 100uL PBS with protease inhibitors. Western blotting for MUC5B was performed as previously described[10,39] with 40uL of supernatant and pellet samples and using a goat polyclonal antibody raised against mouse MUC5B (generously provided by Dr. Alessandra Livraghi-Butrico (UNC)) at 1:500 overnight at 4°C. Samples were randomized by genotype on gel.

*Histology, immunohistochemistry, and colocalization analysis.*
Histological sections were taken from mice sensitized to an immunogenic component of HDM, Der p 1, and challenged with PBS (control) or Der p 1 (allergen challenged) as previously described [10]. Lung tissue was fixed, sectioned, and stained as previously described[10]. Slides were stained with a polyclonal rabbit antibody raised against mouse BPIFB1 (HP8044, Hycult Biotech) at 1:500 and goat polyclonal antibody raised against mouse MUC5B (generously provided by Dr. Alessandra Livraghi-Butrico, UNC) at 1:1000 for 3 hr at room temperature. Slides without primary antibody or with rabbit IgG (JacksonImmuno) were used as controls. Sections incubated with Alexa Fluor 488 anti-rabbit and Alexa Fluor 594 anti-goat secondaries (Invitrogen) at 1:1000 for 1 hr followed by DAPI at 1:1000 for 5 min. Airway cross-sections cut at the hilum/main stem bronchus were selected for imaging. For each animal, 3-5 representative images of airway epithelium containing goblet cells were taken with an Olympus FV1000 confocal microscope (Olympus, Tokyo, Japan) at 60x magnification using uniform settings. Images were analyzed manually using ImageJ (NIH, Bethesda, Maryland). To determine the number of goblet cells per ROI, the total number of airway epithelial cells and goblet cells were counted and calculated as percentage. To establish MUC5B and BPIFB1 colocalization, cells containing MUC5B-positive granules and/or BPIFB1-positive granules were reported as positive for the corresponding marker and calculated as percent of the total number of goblet cells.

*Primary human cell cultures and immune electron microscopy.*
Human bronchial epithelial (HBE) cells were cultured at air-liquid interface (ALI) as previously described.[21] Briefly, HBE cells were seeded on 12 mm diameter Transwell Clear supports (Corning Costar, Cambridge, MA) in Pneumacult EX Plus (StemCell Technologies, Vancouver, Canada). Cultures were maintained at ALI for ~4 weeks in Pneumacult ALI media (StemCell Technologies, Vancouver, Canada) until fully differentiated. HBE cell cultures were fixed in 0.5% glutaraldehyde/4% paraformaldehyde (PFA) in 0.15M sodium phosphate buffer pH 7.4 overnight at 4°C. Samples were washed four times with fixative-free 0.15M sodium phosphate buffer before being incubated overnight at 4°C in blocking solution composed of 0.5M glycine and 1% fish gelatin (FG, VWR, Radnor, PA) in 1x Hanks Balanced Salt Solution (HBSS). Samples were rinsed twice in 1x HBSS before application of a secondary blocking solution of 10% normal donkey serum (MilliporeSigma, Burlington, MA) 1% bovine serum albumin (BSA, Sigma-Aldrich, St. Louis, MO), and 1% FG in HBSS for 20 min at room temperature (RT). Samples were incubated in a primary antibody solution containing either rabbit anti-BPIFB1[40] (generously provided by Dr. Colin Bingle, University of Sheffield) or rabbit anti-MUC5B (H-300/SC-20119, Santa Cruz Biotechnology, Dallas, TX) diluted 1:100 in HBSS/BSA/FG for 1 hour at RT. Negative

controls were run concurrently. Samples were washed four times in HBSS/BSA/FG and incubated in a secondary antibody solution containing biotinylated donkey anti-rabbit IgG (Invitrogen, Carlsbad, CA) diluted 1:200 in 0.05M tris-buffered saline (TBS) and 1% BSA for 30 min at RT. After four washes with TBS/BSA, samples were incubated with streptavidin-conjugated 40nm colloidal gold (nanoComposix, San Diego, CA) diluted 1:75 in TBS/BSA. Samples were washed six times in 0.15M sodium phosphate buffer before secondary fixation in 2% glutaraldehyde/2% PFA for 10 min at RT. Immunolabeled samples were processed for scanning electron microscopy (SEM) by dehydration through increasing concentrations of ethanol followed by critical point drying with liquid $CO_2$. Inserts were cut out and mounted on aluminum SEM stubs using conductive carbon tape and sputter coated with 4nm of chromium. Images were acquired on a Zeiss Supra 25 field emission SEM at a working distance of 8mm and 10kV using backscattered detection.

*Statistics.*

Two-sided Student's *t*-tests or Welch's *t*-tests were used to compare values from WT and KO mice for MCC, CBF, and bioelectric assays. Microbead rheology data were log transformed to and analyzed with an ANCOVA model with genotype and experimental batch as covariates. The statistical framework for RNAseq differential expression was performed according to the DESeq2 vignette. All statistical analyses were carried out in R version 3.6.2. Statistical significance was set at a *P* value of 0.05 or less.

**SUPPLEMENTARY FIGURE LEGENDS**

**Supplementary Figure 1 (Fig S1): Transcriptome-wide differential expression in *Bpifb1* KO mice**. Volcano plot of lung RNAseq differential expression between WT and *Bpifb1* KO mice. Red dots indicate genes with adjusted *P*-value ≤ 0.05 and $|\log_2 (\text{foldchange})| \geq 1$. n=5/genotype. Protein coding genes labeled. Full differential expression data on all genes in Supplementary File 2.

**Supplementary Figure 2 (Fig S2): Airway inflammation in allergen exposed WT and *Bpifb1* KO mice.** Percentage of eosinophils in BAL following house dust mite allergen exposure. n=10/genotype. No difference by genotype in two-sided Welch's *t*-test ($P = 0.2$).

**Supplementary Figure 3 (Fig S3): Additional metrics from microbead rheology and whole BAL imaging.** (A) Fraction of microbeads in sample mucus fraction. (B) Mucus coverage fraction by whole BAL imaging. (C) Size of mucus flakes by whole BAL imaging. (D) Count of mucus flakes by whole BAL imaging. (A-D). No differences by genotype. Boxplots depict median and interquartile ranges. n=10-12/genotype. (E) Relationship between microbead rheology and whole BAL imaging metrics. White dots (WT mice), black dots (*Bpifb1* KO mice). Spearman correlation coefficients on diagonals. Significance of correlation denoted as * $P < 0.05$, ** $P < 0.01$.

**Supplementary Figure 4 (Fig S4): BPIFB1 and MUC5B colocalize in airway epithelial cells of mice without allergen challenge.** Percentage of cells positively stained for BPIFB1 only, MUC5B only, or both in control mice, n=3 mice.

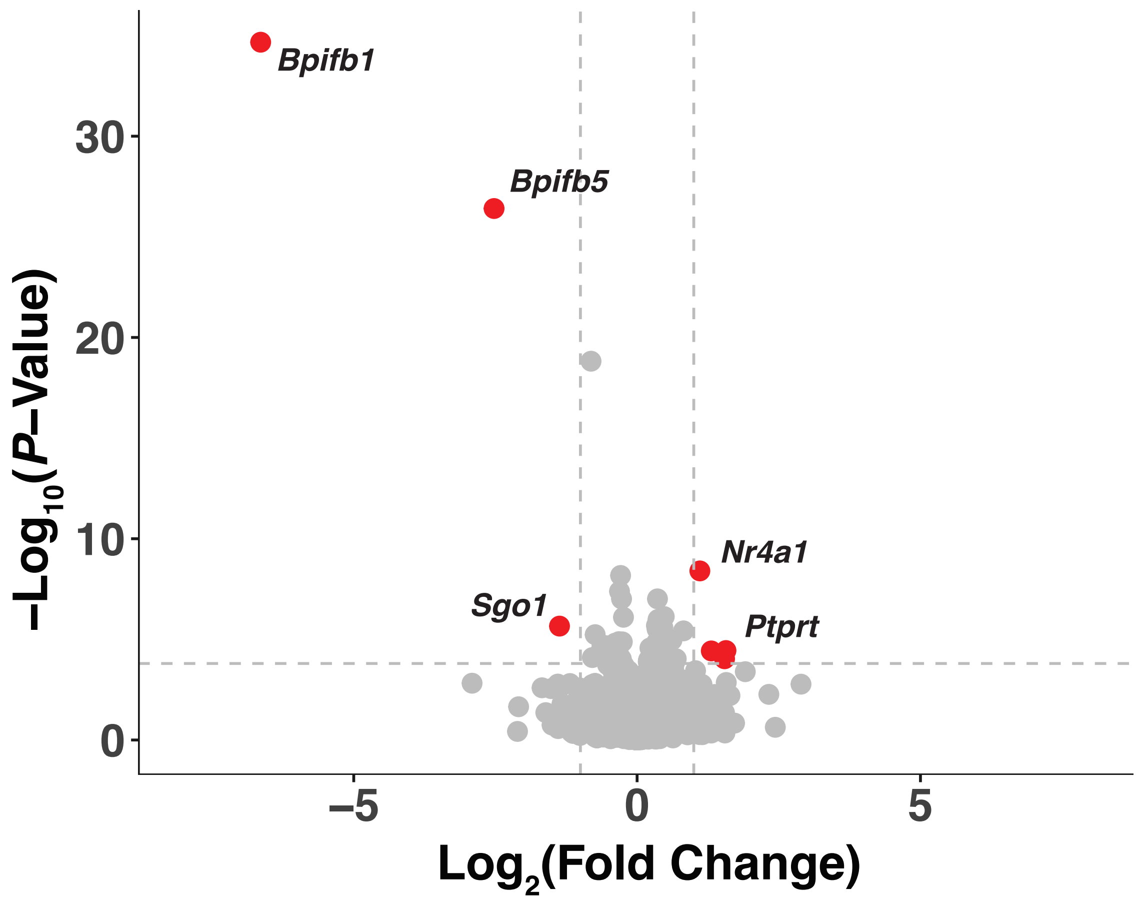


**Supplementary Figure 1 (Fig S1): Transcriptome-wide differential expression in *Bpifb1* KO mice.** Volcano plot of lung RNAseq differential expression between WT and *Bpifb1* KO mice. Red dots indicate genes with adjusted *P*-value ≤ 0.05 and $|\log_2(\text{fold change})| \geq 1$. n=5/genotype. Protein coding genes labeled. Full differential expression data on all genes in Supplementary File 2.

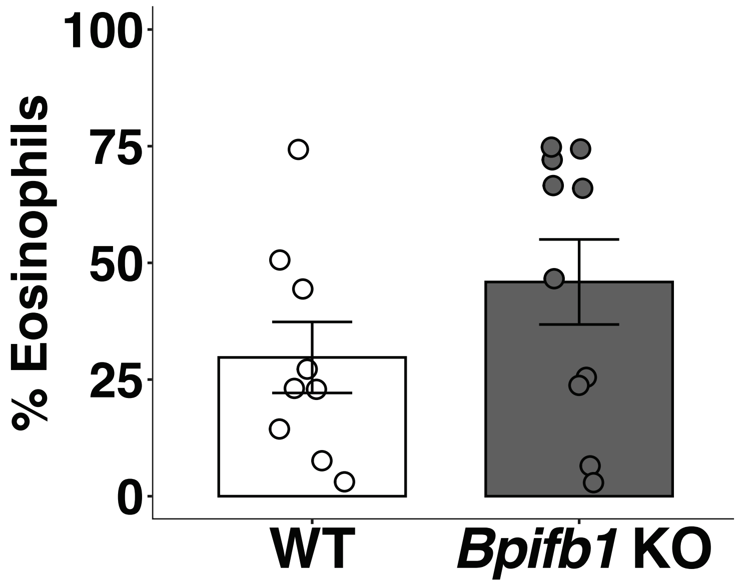


**Supplementary Figure 2 (Fig S2): Airway inflammation in allergen exposed WT and *Bpifb1* KO mice.** Percentage of eosinophils in BAL following house dust mite allergen exposure. n=10/genotype. No difference by genotype in two-sided Welch's *t*-test ($P = 0.2$).

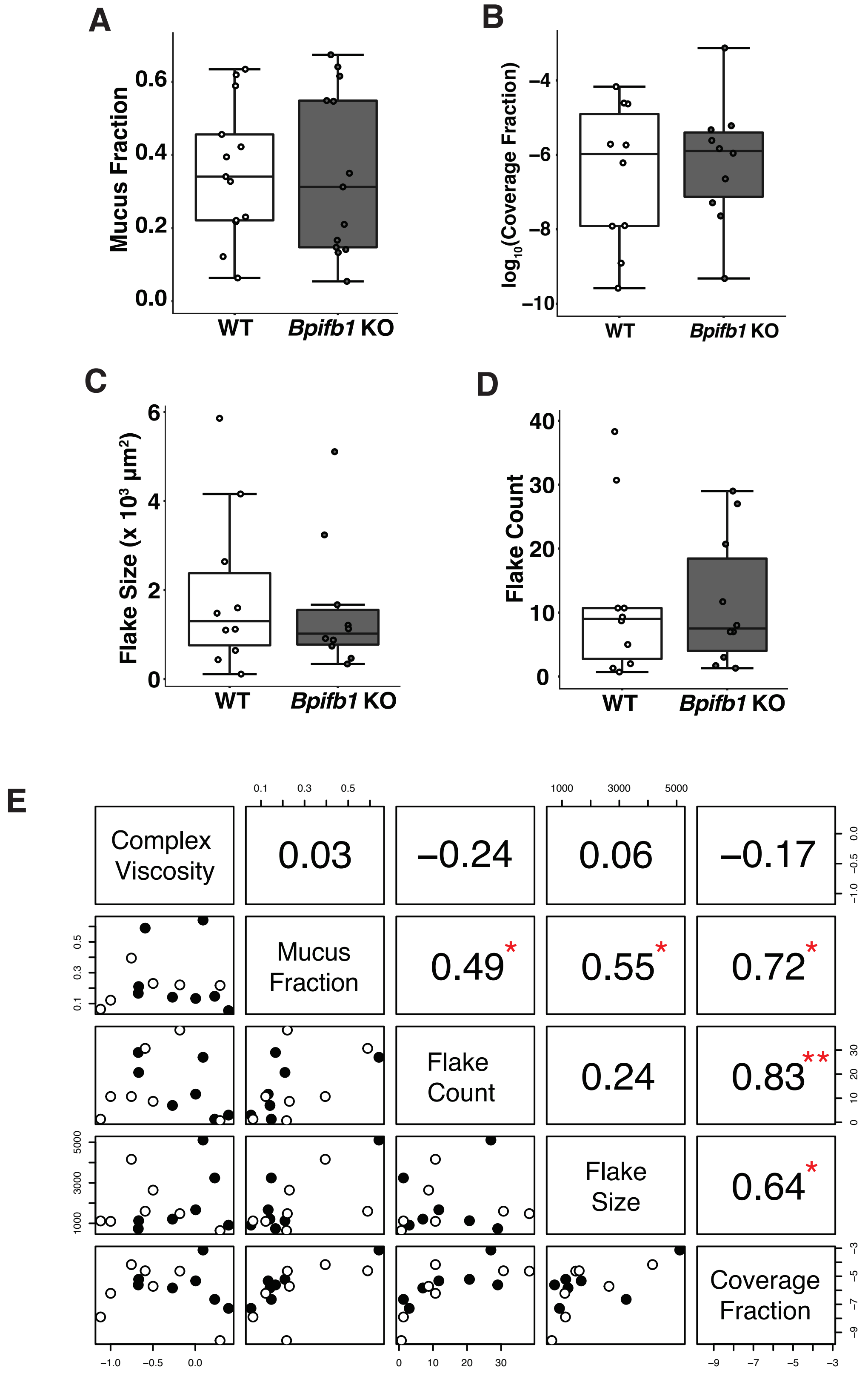


**Supplementary Figure 3 (Fig S3): Additional metrics from microbead rheology and whole BAL imaging.** (A) Fraction of microbeads in sample mucus fraction. (B) Mucus coverage fraction by whole BAL imaging. (C) Size of mucus flakes by whole BAL imaging. (D) Count of mucus flakes by whole BAL imaging. (A-D). No differences by genotype. Boxplots depict median and interquartile ranges. n=10-12/genotype. (E) Relationship between microbead rheology and whole BAL imaging metrics. White dots (WT mice), black dots (*Bpifb1* KO mice). Spearman correlation coefficients on diagonals. Significance of correlation denoted as * $P < 0.05$, ** $P < 0.01$.

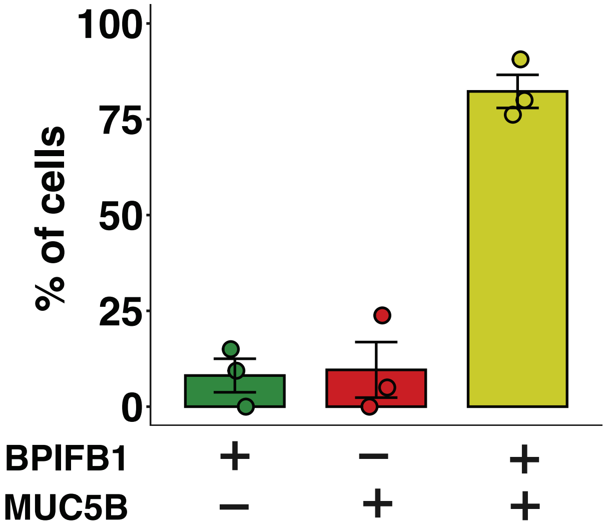


**Supplementary Figure 4 (Fig S4): BPIFB1 and MUC5B colocalize in airway epithelial cells of mice without allergen challenge.** Percentage of cells positively stained for BPIFB1 only, MUC5B only, or both in control mice, n=3 mice.
